# Loss of hepatic phosphoenolpyruvate carboxykinase 1 dysregulates metabolic responses to acute exercise but enhances adaptations to exercise training in mice

**DOI:** 10.1101/2022.09.22.509117

**Authors:** Ferrol I. Rome, Gregory L. Shobert, William C. Voigt, David B. Stagg, Patrycja Puchalska, Shawn C. Burgess, Peter A. Crawford, Curtis C. Hughey

## Abstract

Acute exercise increases liver gluconeogenesis to supply glucose to working muscle. Concurrently, elevated liver lipid breakdown fuels the high energetic cost of gluconeogenesis. This functional coupling between liver gluconeogenesis and lipid oxidation has been proposed to underlie the ability of regular exercise to enhance liver mitochondrial oxidative metabolism and decrease liver steatosis in individuals with non-alcoholic fatty liver disease. Herein we tested whether repeated bouts of increased hepatic gluconeogenesis are necessary for exercise training to lower liver lipids. Experiments used diet-induced obese mice lacking hepatic phosphoenolpyruvate carboxykinase 1 (KO) to inhibit gluconeogenesis and wild type (WT) littermates. ^2^H/^13^C metabolic flux analysis quantified glucose and mitochondrial oxidative fluxes in untrained mice at rest and during acute exercise. Circulating and tissue metabolite levels were determined during sedentary conditions, acute exercise, and refeeding post-exercise. Mice also underwent six weeks of treadmill running protocols to define hepatic and extrahepatic adaptations to exercise training. Untrained KO mice were unable to maintain euglycemia during acute exercise resulting from an inability to increase gluconeogenesis. Liver triacylglycerides were elevated following acute exercise and circulating β-hydroxybutyrate was higher during post-exercise refeeding in untrained KO mice. In contrast, exercise training prevented liver triacylglyceride accumulation in KO mice. This was accompanied by pronounced increases in indices of skeletal muscle mitochondrial oxidative metabolism in KO mice. Together, these results show that hepatic gluconeogenesis is dispensable for exercise training to reduce liver lipids. This may be due to responses in ketone body metabolism and/or metabolic adaptations in skeletal muscle to exercise.

## INTRODUCTION

An outcome of society’s modernization is that individuals lead sedentary lifestyles for which they are poorly adapted. Lower physical activity is associated with non-alcoholic fatty liver disease (NAFLD), which is a spectrum of liver pathologies ranging from hepatic steatosis to non-alcoholic steatohepatitis that afflicts approximately 30% of the global population (1-5). The ectopic lipid deposition in the liver increases the risk for advanced liver diseases (cirrhosis and hepatocellular carcinoma) and is associated with a cluster of diseases comprising the metabolic syndrome (3, 6, 7). Currently, there are no approved pharmacotherapies for NAFLD, however, prior research has shown that regular physical activity or exercise reduces hepatic steatosis through both hepatic and extrahepatic mechanisms (8-12). At the level of the liver, regular exercise diminishes hepatic lipid content partly by enhancing the disposal of lipids in mitochondrial oxidative metabolism pathways (9, 13). More specifically, exercise training increases the molecular machinery and/or functional capacity of liver β-oxidation, tricarboxylic acid (TCA) cycle flux, and oxidative phosphorylation (9, 13).

Gluconeogenesis is energetically linked to liver mitochondrial oxidative metabolism during exercise. To maintain glucose homeostasis during acute exercise, the liver increases glucose production to match the heightened rate of muscle glucose uptake needed to meet the energy demands of muscular work (14, 15). Elevated glucose production is mediated through hepatic glycogenolysis and by channeling intermediates such as glycerol and phosphoenolpyruvate (via pyruvate, lactate, and amino acids) into the gluconeogenic pathway (14-18). Importantly, gluconeogenesis is energetically costly, necessitating its functional coupling to mitochondrial fat oxidation to meet the high ATP requirements (9, 13, 18, 19). Thus, hepatic gluconeogenesis provoked by muscular work may lower liver lipids by promoting lipid oxidation. Support for this mechanism comes from studies in mice with impaired activity of phosphoenolpyruvate carboxykinase 1 (PCK1), which generates phosphoenolpyruvate from oxaloacetate and allows TCA cycle intermediates to be converted to glucose. Global knockdown or liver-specific deletion of PCK1 impairs gluconeogenesis from phosphoenolpyruvate, suppresses hepatic TCA cycle fluxes, and elevates liver triacylglycerides under fasted, sedentary conditions (20-24).

The work presented herein tested the hypothesis that repeated bouts of acutely increased hepatic gluconeogenesis are necessary for exercise training to lower liver lipids. Experiments leveraged diet-induced obese mice with a liver-specific deletion of PCK1 (KO) and wild type (WT) littermates. Stable isotope infusions in conscious mice, ^2^H/^13^C metabolic flux analysis, and targeted metabolomics were performed to determine endogenous glucose and associated mitochondrial oxidative fluxes at rest and during an acute treadmill running bout in untrained mice. WT and KO mice also underwent six weeks of treadmill running protocols to define hepatic and extrahepatic adaptations to exercise training. We report that, surprisingly, hepatic gluconeogenesis is dispensable for exercise training to reduce liver lipids.

## MATERIALS AND METHODS

### Mouse Models and Husbandry

The University of Minnesota Institutional Animal Care and Use Committee approved all procedures performed in mice. Mice on a C57BL/6J background with a hepatic-specific PCK1 KO were generated by breeding mice expressing Cre recombinase under control of the albumin promoter (B6.Cg-Tg(Alb-cre)21Mgn/J) with PCK1 floxed mice as previously described (20). Floxed PCK1 littermates of KO mice were used as WT controls. Mice received Teklad Global

18% Protein Rodent Diet (2918, Madison, WI) from weaning at three weeks of age until six weeks of age. At six weeks of age, all mice were provided a high-fat diet (60% kcal from fat, D12492; Research Diets Inc., New Brunswick, NJ). Food and water were provided ad libitum to mice housed with cellulose bedding (Cellu-nest™, Shepherd Specialty Papers, Watertown, TN) in temperature- and humidity-controlled conditions maintained on a 14:10 hour light/dark cycle. Male mice were used for all experiments and were assigned to one of four experimental cohorts (See Fig. 1A for a schematic representation of the experimental cohort protocols and timeline).

**FIGURE 1.**
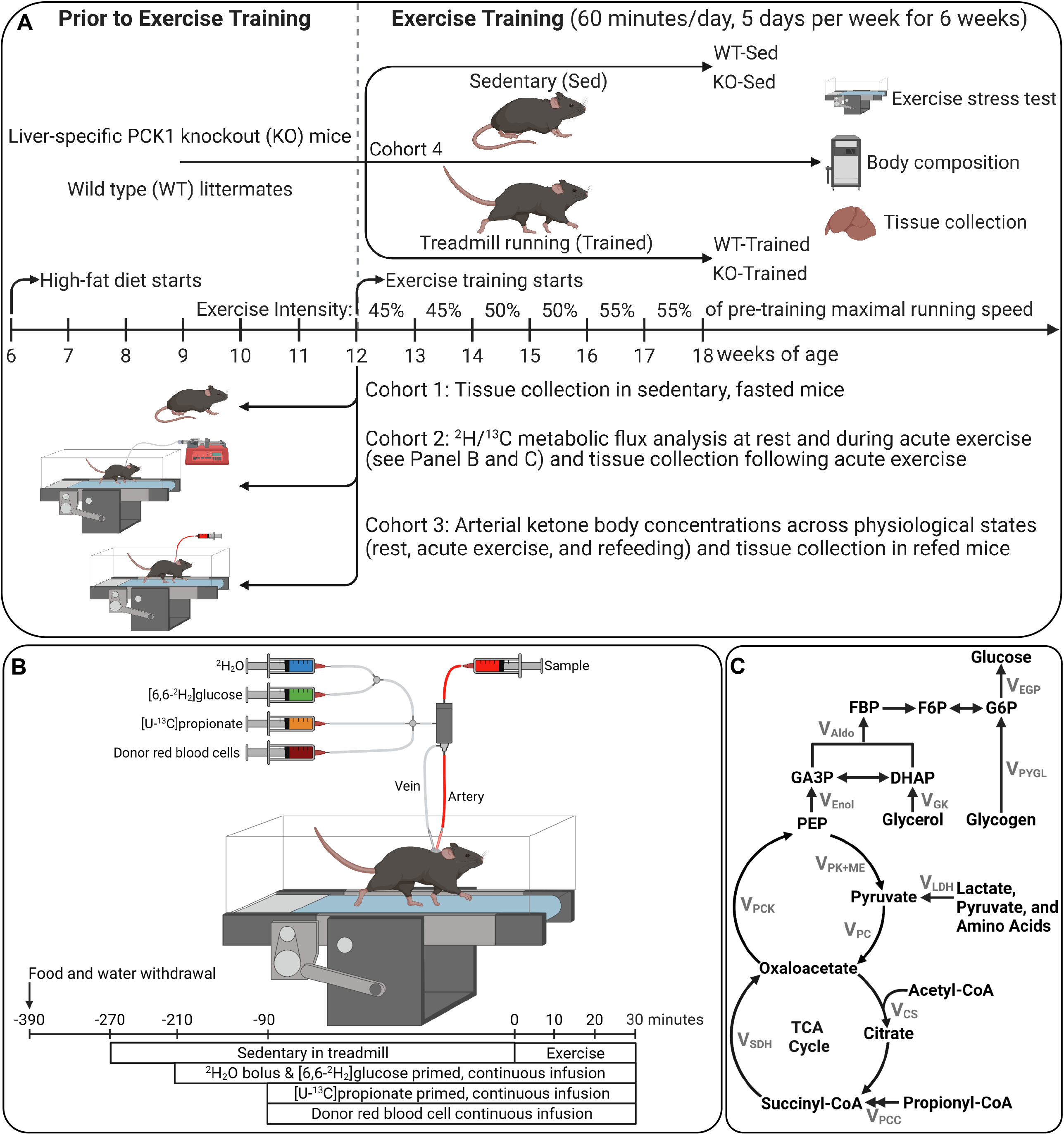
Schematic representation of experimental timeline and procedures. **(A)** Mice with a liver-specific deletion of phosphoenolpyruvate carboxykinase 1 (KO) and wild type (WT) littermates were provided a high-fat diet starting at 6 weeks of age and assigned to four experimental cohorts. Cohort 1 mice had tissue collected at 12 weeks of age under fasted, sedentary conditions. Cohort 2 mice had vascular catheters implanted at 11 weeks of age and received stable isotope infusions to quantify endogenous glucose and associated oxidative fluxes during rest and acute exercise at 12 weeks of age. Cohort 3 mice had an arterial catheter implanted at 11 weeks of age for ketone body measurements under sedentary conditions, acute exercise, post-exercise fasting, and post-exercise refeeding at 12 weeks of age. Cohort 4 mice underwent exercise training protocol consisting of treadmill running 60 minutes a day, five days per week for 6 weeks starting at 12 weeks of age. Pre-and post-exercise training analyses included body composition and exercise stress tests to evaluate maximal running speed. Tissues were collected from all mice upon completion of *in vivo* experiments for molecular analysis. **(B)** Stable isotope infusions at rest and during acute treadmill running bout were performed in mice 7 days following carotid arterial and jugular catheter implantation surgeries. At 210 minutes prior to the treadmill running bout (3 hours of fasting), a ^2^H_2_O bolus was administered into the venous circulation to enrich total body water at 4.5%. A [6,6-^2^H_2_]glucose prime was infused followed by a continuous infusion was initiated with the ^2^H_2_O bolus. Ninety minutes before the onset of exercise (5 hours of fasting), a primed, continuous infusion of [U-^13^C]propionate was started. Donor red blood cells were administered to prevent a decline in hematocrit. Arterial samples were obtained prior to stable isotope infusion as well as during 30-minute exercise bout for ^2^H/^13^C metabolic flux analysis. **(C)** Schematic representation of glucose and associated nutrient fluxes quantified by ^2^H/^13^C metabolic flux analysis.

Cohort 1 mice. At 12 weeks of age, mice were fasted under sedentary conditions for 7 hours (starting within the first hour of the light cycle) and euthanized via cervical dislocation. Tissues were rapidly excised, freeze-clamped in liquid nitrogen, and stored at -80°C.

Cohort 2 mice. At 11 weeks of age, catheters were implanted in the jugular vein and carotid artery for isotope infusion and sampling protocols as previously described (25, 26). The exteriorized ends of the implanted catheters were flushed with 200 U·mL^-1^ heparinized saline and sealed with stainless-steel plugs. Following surgery, mice were housed individually and provided 7-8 days of post-operative recovery prior to stable isotope infusions studies during rest and acute exercise (See Stable isotope infusions).

Cohort 3 mice. At 11 weeks of age, a catheter was placed in the carotid artery. Mice were individually housed and provided 7-8 days of post-surgical recovery prior to sampling protocols to obtain blood β-hydroxybutyrate (βOHB) measurements during sedentary conditions, acute exercise, and refeeding (See Serial β-ketone body measurements).

Cohort 4 mice. At 12 weeks of age, body composition was measured and the exercise training protocols were initiated. Mice completed a 60-minute treadmill running bout, five days a week for six weeks. The treadmill running speed was 45% of the mouse’s baseline maximal running speed during the first two weeks of the exercise training protocol and increased progressively by 5% of the maximal running speed every two weeks. Sedentary mice were placed in an enclosed container on top of the treadmill. Similar to the exercise-trained mice, this intervention was 60 minutes per day, 5 days per week for six weeks. Following the exercise training protocols, body composition was re-assessed, and maximal running speed was re-evaluated via an exercise stress test. Forty-eight hours after the exercise stress test mice were fasted for 7 hours under sedentary conditions (starting within the first hour of the light cycle) and euthanized via cervical dislocation. Tissues were rapidly excised, freeze-clamped in liquid nitrogen, and stored at -80°C.

### Exercise Stress Test

Mice underwent two exercise stress tests to determine maximal running speed. The initial exercise stress test was completed 24-48 hours before the start of the 6-week exercise training protocol. The second exercise stress test was executed 24-48 hours following the completion of the 6-week exercise training protocol. The exercise stress test protocols were performed as previously described (18, 27). Briefly, a 10-minute exercise bout at 10 m·min^−1^ (0% incline) was completed 24 hours prior to the exercise stress test to acclimate mice to running on an enclosed single lane treadmill (Columbus Instruments, Columbus, OH, US). For the exercise stress test, mice were placed in the same enclosed single lane treadmill and the initial treadmill belt speed was 10 m·min^-1^ (0% incline). The treadmill belt speed was increased by 4 m·min^-1^ at 3-minute intervals until exhaustion. Exhaustion was defined as the point in which the mouse remained on the shock grid at the posterior of the treadmill for more than five continuous seconds.

### Body Composition

The initial body composition measurement was obtained using an EchoMRI-100™ Body Composition Analyzer (EchoMRI LLC, Houston, TX) in 12-week-old mice before the first acclimation exercise bout of the exercise stress test procedures. The second body composition measurements were acquired in 18-week-old mice prior to the final 60-minute treadmill running bout.

### Stable isotope infusions

All mice were within 10% of pre-surgical weight prior to stable isotope infusions. During the first hour of the housing light cycle, both food and water were restricted for the entirety of the experiment (7 hours). Two hours into the fast, mice were placed in the enclosed single lane treadmill and the exteriorized catheters were connected to infusion syringes. Following a one-hour acclimation period, an 80 μl arterial blood sample was taken to determine natural isotopic enrichment of plasma glucose. Immediately after this sample acquisition, a stable isotope infusion protocol was initiated to allow for the quantification of glucose and associated oxidative fluxes as previously performed (18). Briefly, a ^2^H_2_O (99.9%)-saline bolus containing [6,6- ^2^H_2_]glucose (99%) was infused over 25 minutes to both enrich the H_2_O pool and administer a [6,6-^2^H_2_]glucose prime (440 μmol·kg^-1^). A continuous infusion of [6,6-^2^H_2_]glucose (4.4 μmol·kg^-1^·min^-1^) was initiated after the ^2^H_2_O-saline bolus and [6,6-^2^H_2_]glucose prime. A primed (1.1 mmol·kg^-1^), continuous (0.055 mmol·kg^-1^·min^-1^) intravenous infusion of [U-^13^C]propionate was started two hours after the ^2^H_2_O bolus and [6,6-^2^H_2_]glucose prime. Four, 100 µl arterial samples were acquired 90-120 minutes following the [U-^13^C]propionate bolus (time = 0-30 minutes of treadmill running) to determine arterial glucose concentration and enrichment. The sample obtained at 90 minutes following the [U-^13^C]propionate bolus (time = 0 minutes of treadmill running) was obtained while mice were in a sedentary state on a stationary treadmill. Samples taken 100-120 minutes following the [U-^13^C]propionate bolus (time = 10-30 minutes of exercise) were obtained while mice were completing an acute treadmill running bout at 12 m·min^-1^. All plasma samples were stored at -80°C. Donor erythrocytes were provided by constant rate infusion throughout the experiment to prevent a decline in hematocrit. Mice were removed from the treadmill and sacrificed by cervical dislocation following the final sample. Tissues were rapidly excised, freeze-clamped in liquid nitrogen, and stored at -80°C. A schematic of the stable isotope infusion and acute exercise protocols is provided in Fig. 1B. All stable isotopes were purchased from Cambridge Isotope Laboratories, Inc. (Tewksbury, MA).

### Glucose Derivatization and GC-MS Analysis

Approximately 40 µl of plasma obtained prior to the stable isotope infusion and at the 0, 10, 20, and 30 minute time points of the isotope infusions were used for di-*O*-isopropylidene propionate, aldonitrile pentapropionate, and methyloxime pentapropionate derivatives of glucose (18). GC-MS analysis was performed and uncorrected mass isotopomer distributions (MIDs) for six fragment ions were determined as previously described (26).

### ^2^H/^13^C Metabolic Flux Analysis

The metabolic flux analysis methodology employed in these studies followed that previously described with minor modifications (26, 28). To summarize, a reaction network (29) was constructed using Isotopomer Network Compartmental Analysis (INCA) software (30). The reaction network defined the carbon and hydrogen transitions for hepatic glucose and associated oxidative metabolism reactions. The flux through each network reaction was determined relative to citrate synthase flux (V_CS_) by minimizing the sum of squared residuals between simulated and experimentally determined MIDs of the six fragment ions previously described. Flux estimates were repeated 50 times from random initial values. Goodness of fit was assessed by a chi-square test (*p* = 0.0*5*) and confidence intervals of 95% were determined as previously described (28). Mouse body weights and the [6,6-^2^H_2_]glucose infusion rate were used to determine absolute values.

### Serial β-ketone body measurements

All mice were within 10% of pre-surgical weight prior to initiating the experiments. Both food and water were restricted within the first hour of the housing light cycle. Two hours into the fast, mice were placed in the enclosed single lane treadmill and the exteriorized arterial catheter were connected a sampling swivel. Following a one-hour acclimation period (3 hours of fasting), an arterial blood sample was taken to determine blood βOHB with a Precision Xtra Blood Glucose & Ketone meter (Abbott Diabetes Care, Alameda, CA). Three additional βOHB measurements were taken with mice on a stationary treadmill at 5, 6, and 6.5 hours of fasting. Mice then underwent a 30-minute treadmill running bout at 12 m·min^-1^ wherein βOHB levels were determined every 10 minutes. Subsequently, mice remained fasting on a stationary treadmill to access post-exercise ketosis every 10 minutes for 90 minutes. For the final hour of the experiment, mice were placed in a container with bedding and ad libitum access to food. βOHB was assessed during this refeeding phase every 15 minutes. Mice were then euthanized via cervical dislocation and tissues were rapidly excised, freeze-clamped in liquid nitrogen, and stored at -80°C.

### Hormone and metabolite analyses

Blood glucose was measured with a Contour® blood glucose meter (Ascensia Diabetes Care, Parsippany, NJ). Plasma non-esterified fatty acids (NEFAs) were quantified via the Wako HR series NEFA-HR(2) assay (FUJIFILM Medical Systems USA, Lexington, MA). Plasma insulin was determined using the Mercodia Ultrasensitive Mouse Insulin ELISA (Winston Salem, NC). Plasma and liver acetoacetate (AcAc), βOHB, and total ketone bodies (TKB) were quantified via UPLC-MS/MS as previously detailed (31). Liver, kidney, and gastrocnemius glycogen were quantified as previously described (26, 32). Liver phospholipids (PL), diacylglycerides (DAG), and triacylglycerides (TAG) during sedentary conditions were evaluated as previously outlined (26, 33). Liver TAG following acute exercise and during refeeding were assessed with the Triglycerides - Liquid Reagent Set (Pointe Scientific, Inc., Lincoln Park, MI).

### Immunoblotting

Tissue homogenates were prepared as previously detailed (18, 26, 34, 35). Tissue proteins (15 µg) were separated using electrophoresis on a NuPAGE 4-12% Bis-Tris gel (Invitrogen, Carlsbad, CA, USA) and transferred to a PVDF membrane. Immunoblotting was performed with the following primary antibodies: β-hydroxybutyrate dehydrogenase (BDH1; Proteintech, Rosemont, IL, USA, 15417-1-AP; RRID:AB_2274683), fructose 1,6-bisphosphatase (FBP; Proteintech; 12842-1-AP; RRID:AB_2103572), glucose-6-phosphatase (G6PC; Proteintech, 22169-1-AP; RRID:AB_2879015), glycogen phosphorylase (PYGL; Proteintech, 15851-1-AP; RRID:AB_2175014), phospho-glycogen synthase (Ser641) (pGS^S641^; Cell Signaling Technology, 3891; RRID:AB_2116390), glycogen synthase 2 (GS; Proteintech, 22371-1-AP; RRID:AB_2879091), mitochondrial encoded cytochrome c oxidase II (MT-CO2, Proteintech, 55070-1-AP; RRID:AB_10859832), NADH dehydrogenase (ubiquinone) iron-sulfur protein 4 (NDUFS4, Abcam, ab139178; RRID:AB_2922810), cytosolic phosphoenolpyruvate carboxykinase (PCK1; Proteintech, 16754-1-AP; RRID:AB_2160031), succinyl-CoA:3-oxoacid-CoA transferase (SCOT, Proteintech, 12175-1-AP; RRID:AB_2157444), Total OXPHOS Rodent WB Antibody Cocktail (Abcam, Cambridge, MA, USA, ab110413; RRID:AB_2629281), and 3-hydroxymethylglutaryl-CoA synthase 2 (HMGCS2; Cell Signaling Technology, 20940S; RRID:AB_2798853). All primary antibodies were used as a dilution of 1:1000. The PVDF membranes were treated with a chemiluminescent substrate (ThermoFisherScientific, Waltham, MA, USA) and imaged using a ChemiDoc™ Imaging system and Image Lab™ software (Bio-Rad, Hercules, CA, USA). Total protein was assessed via BLOT-FastStain (G-Bioscience, St. Louis, MO, USA) and used as the loading control. Densitometry was completed using ImageJ software.

### Statistical analyses

GraphPad Prism software (GraphPad Software LLC., San Diego, CA) was used to perform Student’s t test, two-way ANOVAs followed by Tukey’s post-hoc tests, and repeated measures ANOVAs followed by Šidák’s post-hoc tests as appropriate to detect statistical differences (p<0.05). All data are reported as mean ± SEM.

## RESULTS

### An inability to increase gluconeogenesis prevents euglycemia in KO mice during exercise

At 12 weeks of age, plasma insulin (Fig. 2A) and NEFAs (Fig. 2B) were comparable between genotypes prior to and during a 30-minute treadmill run. Blood glucose was similar between genotypes at rest (Fig. 2C). In contrast, circulating glucose was lower in KO mice compared to WT mice during exercise (Fig. 2C). Stable isotope infusions combined with ^2^H/^13^C metabolic flux analysis allowed glucose and associated oxidative fluxes to be quantified (Fig. 1C and Fig. 2D-O). Endogenous glucose production (V_EGP_) was comparable between genotypes at rest but reduced in KO mice compared to WT mice during exercise (Fig. 2D). Glycogenolysis (V_PYGL_) was comparable between WT and KO mice prior to and during exercise (Fig. 2E).

**FIGURE 2.**
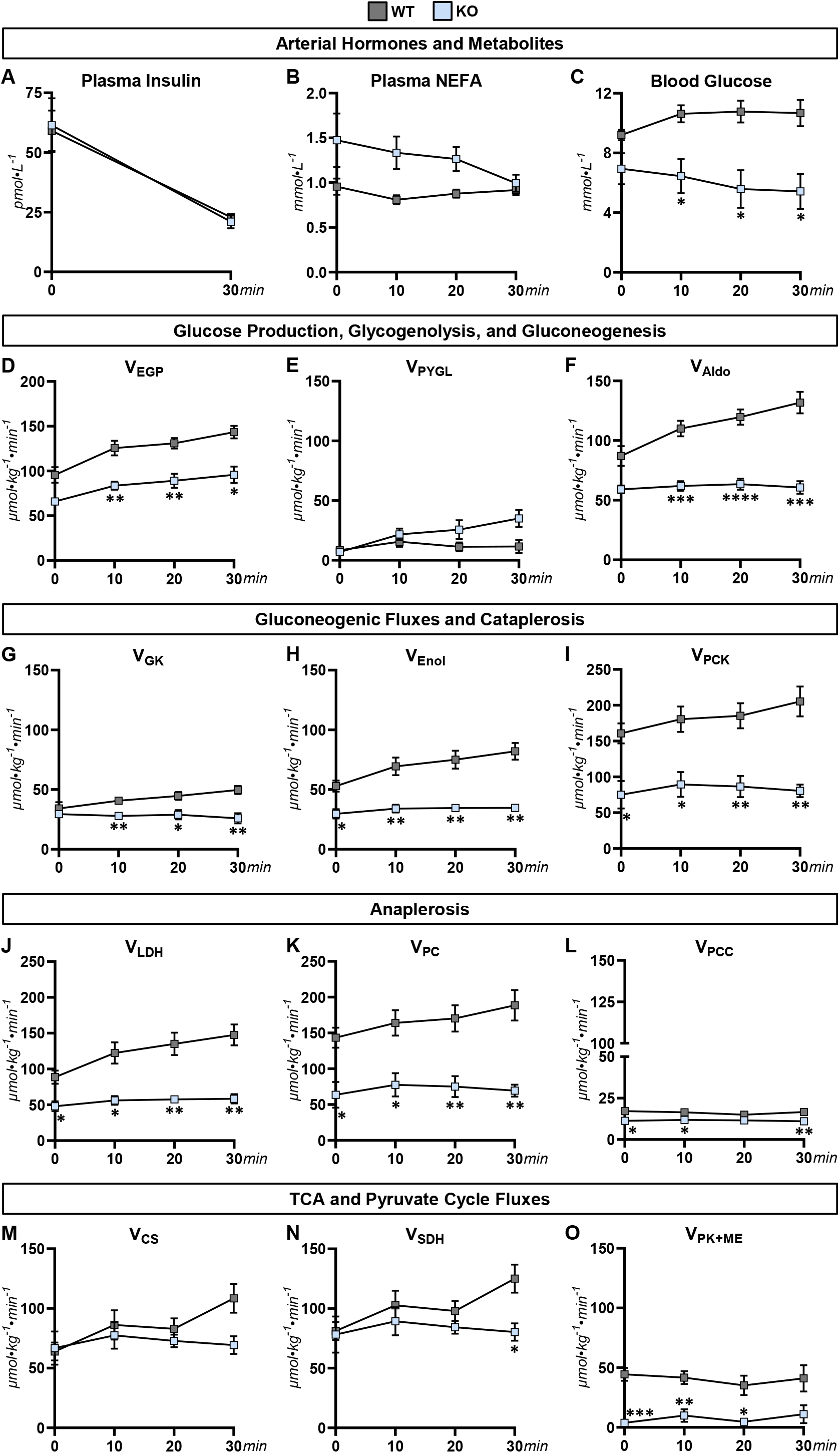
Glucose producing and oxidative metabolism fluxes prior to and during treadmill running in mice lacking liver PCK1. **(A)** Plasma insulin (pmol·L^-1^) prior to exercise (0-minute time point) and at the conclusion of a 30-minute treadmill run (30-minute time point) in 12-week-old mice with a liver-specific deletion of PCK1 (KO) and wild type (WT) littermates. A time course of **(B)** plasma NEFA and **(C)** blood glucose concentration (mmol·L^-1^) prior to and during a 30-minute treadmill run. Model-estimated, nutrient fluxes (µmol·kg^-1^·min^-1^) in WT and KO mice prior to and during a 30-minute of treadmill run for **(D)** endogenous glucose production (V_EGP_), **(E)** glycogenolysis (V_PYGL_), **(F)** total gluconeogenesis (V_Aldo_), **(G)** gluconeogenesis from glycerol (V_GK_), **(H)** gluconeogenesis from phosphoenolpyruvate (V_Enol_), **(I)** tricarboxylic acid cycle cataplerosis (V_PCK_), **(J)** flux from unlabeled, non-phosphoenolpyruvate, anaplerotic sources to pyruvate (V_LDH_), **(K)** anaplerosis from pyruvate (V_PC_), **(L)** anaplerosis from propionyl-CoA (V_PCC_), **(M)** flux from oxaloacetate and acetyl-CoA to citrate (V_CS_), **(N)** flux from succinyl-CoA to oxaloacetate (V_SDH_), and **(O)** pyruvate cycling (V_*PK+ME*_). n = 5-7 per genotype. Data are mean ± SEM. *p<0.05, **p<0.01, ***p<0.001, and ****p<0.001 vs. WT at specified time point.

Similar to V_EGP_, total gluconeogenesis (V_Aldo_; Fig. 2F) and gluconeogenesis from glycerol (V_GK_; Fig. 2G) were comparable between genotypes under sedentary conditions, but lower in KO mice during exercise. Gluconeogenic flux from phosphoenolpyruvate (V_Enol_; Fig. 2H), cataplerosis (V_PCK_; Fig. 2I), and anaplerotic fluxes (V_LDH_, V_PC_, and V_PCC_; Fig. 2J-L) were blunted in KO mice compared to WT mice at rest and during exercise. TCA cycle fluxes (V_CS_ and V_SDH_) were largely comparable between genotypes, but V_SDH_ was higher in WT mice at 30 minutes of treadmill running (Fig. 2M and N). Pyruvate cycling (V_PK+ME_) was suppressed in KO mice compared to WT mice at rest and during exercise (Fig. 2O). These results show that the inability of muscular work to stimulate gluconeogenic fluxes in KO mice above resting rates compromises glucose production and homeostasis during acute exercise.

### Loss of hepatic PCK1 compromises glycogen pools in liver and muscle

In conjunction with glucose fluxes, mediators of glucose production and whole-body glucose control were assessed in liver, muscle, and kidney tissue of WT and KO mice (Fig. 3). Glycogen phosphorylase (PYGL) and the phospho-glycogen synthase-to-glycogen synthase ratio (pGS^S641^/GS) were decreased and increased, respectively, in the livers of KO mice compared to WT mice (Fig. 3A). The decline in these regulators of glycogen availability was accompanied by lower liver glycogen in KO mice at rest (Fig. 3B), following acute exercise (Fig. 3C), and in the refed state following acute exercise (Fig. 3D). Perturbations in glycogen metabolism were not confined to the livers of KO mice. Skeletal muscle glycogen was comparable between WT and KO mice under fasted, sedentary conditions (Fig. 3E), but, consistent with the hypoglycemic phenotype during exercise, muscle glycogen trended lower in KO mice following a 30-minute treadmill run (Fig. 3F) and in response to refeeding following acute exercise (Fig. 3G). Kidney glycogen was not detectable during fasting (Fig. 3H) or following exercise and refeeding (data not shown) in either WT or KO mice. However, the gluconeogenic enzyme fructose 1,6-bisphosphatase (FBP) was increased in kidney tissue of KO mice compared to WT mice (Fig. 3I), which is consistent with prior work showing renal gluconeogenesis to be elevated in mice lacking hepatic PCK1 (24). Our results highlight extensive remodeling of both hepatic and extrahepatic glucose metabolism occurs in response to the loss of hepatic PCK1 despite preserved euglycemia at rest.

**FIGURE 3.**
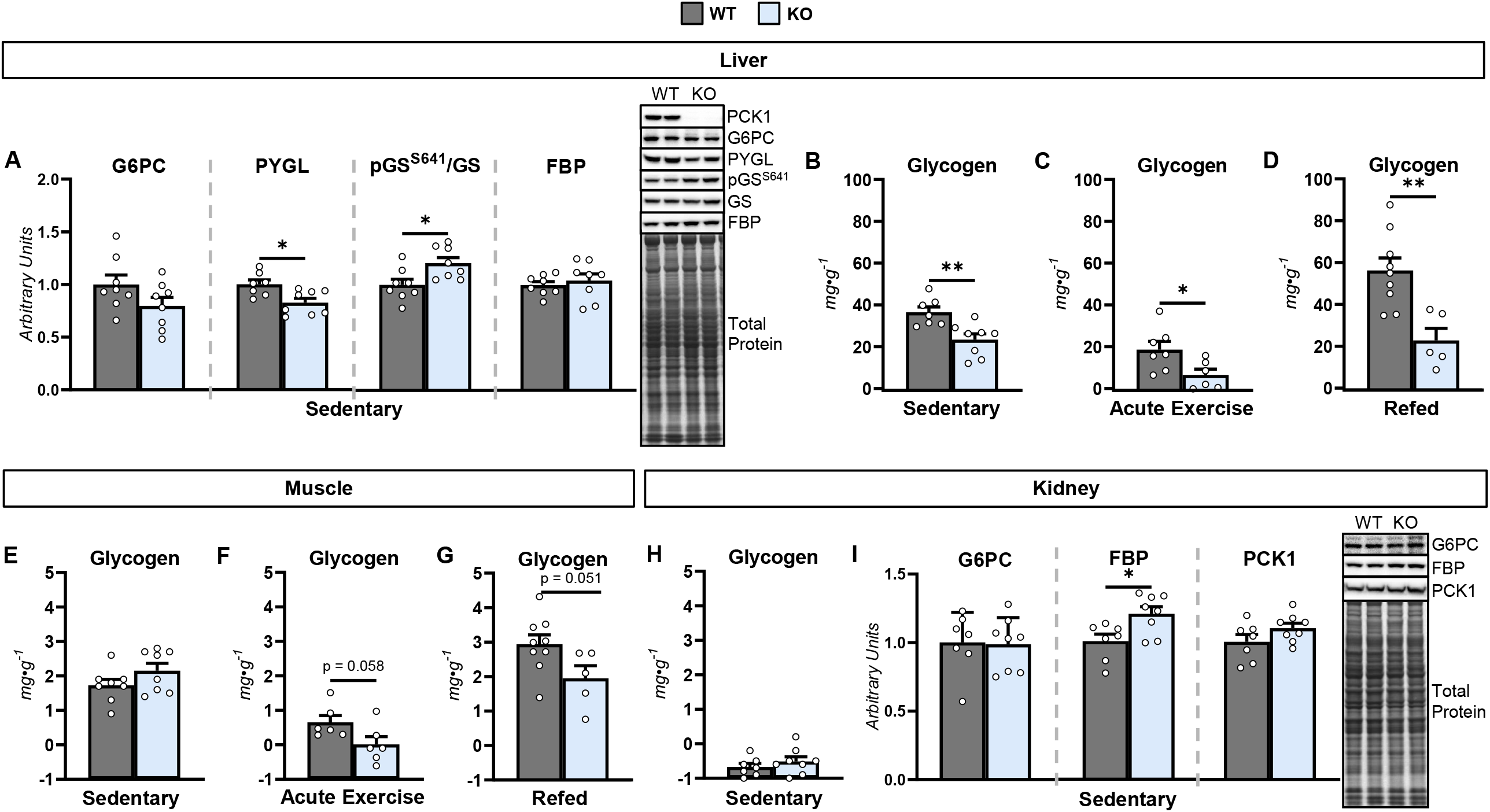
Tissue glucose and glycogen homeostasis in mice lacking liver PCK1. Livers from 12-week-old mice with a hepatic-specific knockout (KO) of phosphoenolpyruvate carboxykinase 1 (PCK1) and wild type (WT) littermates were obtained during fasted, sedentary mice to determine **(A)** liver PCK1, glucose-6-phosphatase catalytic subunit (G6PC), glycogen phosphorylase (PYGL), phosphorylated glycogen synthase-to-total glycogen synthase ratio (pGS^S641^/GS), and fructose 1,6-biphosphatase (FBP) protein via immunoblotting. Liver glycogen (mg·g^-1^) in **(B)** sedentary mice prior to acute exercise, **(C)** mice following 30-minutes of exercise, and **(D)** mice refed following acute exercise. Muscle glycogen (mg·g^-1^) in **(E)** sedentary mice prior to acute exercise, **(F)** mice following 30-minutes of exercise, and **(G)** mice refed following acute exercise. **(H)** Kidney glycogen (mg·g^-1^) in sedentary mice. **(I)** Kidney G6PC, FBP, and PCK1 as determined by immunoblotting and representative immunoblots. n = 5-9 per genotype. Data are mean ± SEM. *p<0.05 and **p<0.01.

### Ketone body dynamics are dysregulated in KO mice

In addition to glucose, ketone bodies are predominantly liver-derived metabolites that provide an additional source of energy for extrahepatic tissues under conditions of energetic stress such as fasting and exercise. The protein expression of ketogenic enzymes, 3-hydroxymethylglutaryl-CoA synthase 2 (HMGCS2) and β-hydroxybutyrate dehydrogenase 1 (BDH1), was similar between WT and KO mice (Fig. 4A). Plasma acetoacetate (AcAc) was decreased (Fig. 4B) and β-hydroxybutyrate (βOHB) was unchanged in KO mice compared to WT mice at rest and following a 30-minute treadmill run (Fig. 4C). Thus, there was a 3-4-fold increase in the βOHB-to-AcAc ratio in KO mice (Fig. 4D), suggesting markedly reduced redox state in liver as previously described (36). To further evaluate ketone body metabolism, circulating βOHB was measured across multiple physiological states including pre-exercise sedentary conditions, acute exercise, post-exercise sedentary conditions, and in response to refeeding. Notably, blood βOHB was elevated during the refed state following exercise in KO mice compared to WT mice (Fig. 4E and F). Since β-oxidation supplies reducing equivalents for mitochondrial oxidative phosphorylation, respiratory chain complexes were examined. Liver mitochondrial respiratory complexes (CI-CV) were comparable between genotypes under fasted, sedentary conditions (Fig. 4G). Liver lipids were examined given that ketogenesis requires an ample supply of fatty acid delivery. Liver phospholipids (PL), diacylglycerides (DAG), and triacylglycerides (TAG) were unchanged by the loss of hepatic PCK1 at rest (Fig. 4H-J).

**FIGURE 4.**
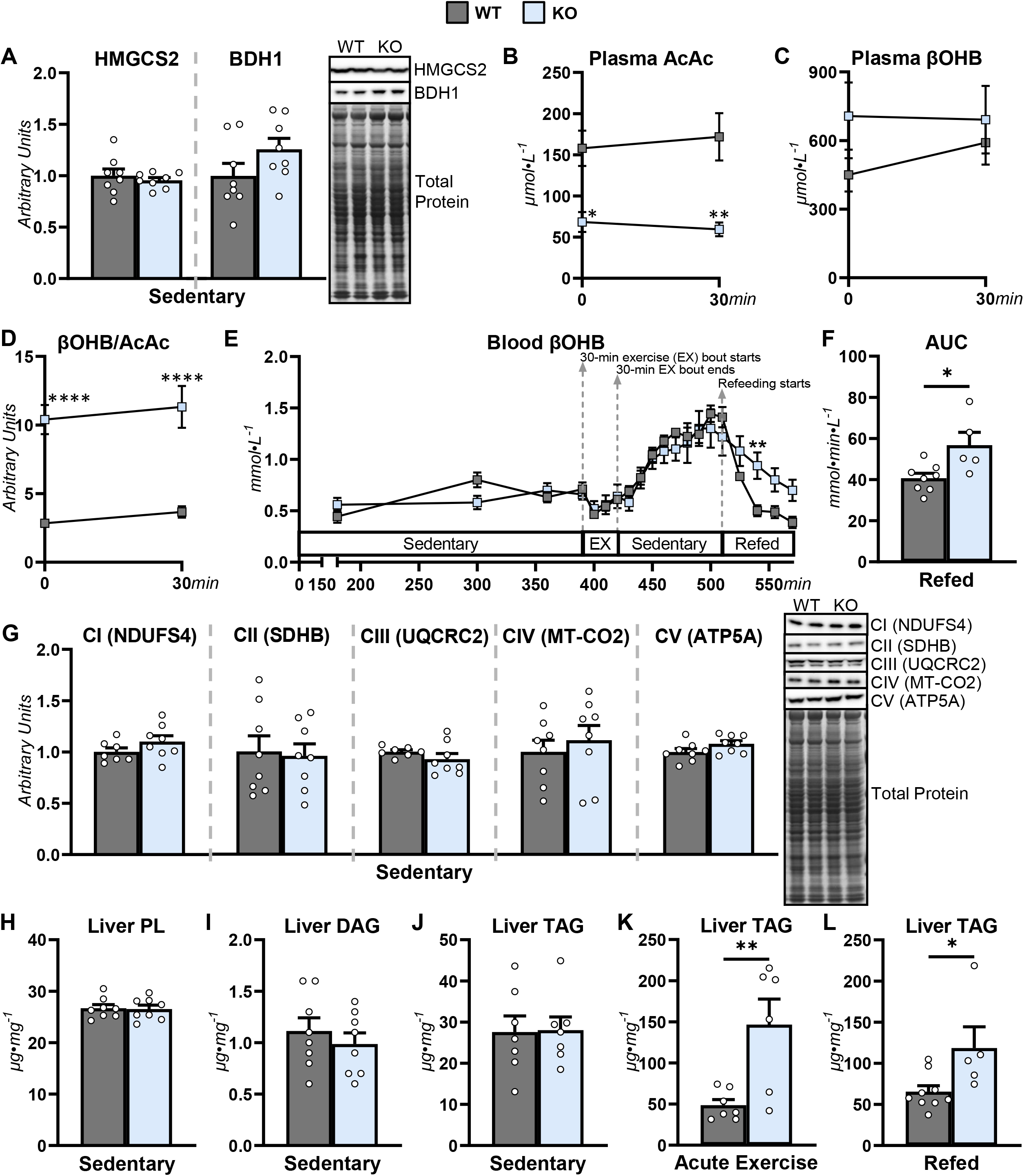
Ketone body and lipid homeostasis in mice lacking liver PCK1. Livers from 12-week-old mice with a hepatic-specific knockout (KO) of phosphoenolpyruvate carboxykinase 1 (PCK1) and wild type (WT) littermates were obtained during fasted, sedentary conditions to assess **(A)** liver 3-hydroxymethylglutaryl-CoA synthase 2 (HMGCS2) and β-hydroxybutyrate dehydrogenase (BDH1) via determined by immunoblotting. Plasma **(B)** acetoacetate (AcAc; µmol·L^-1^), **(C)** β-hydroxybutyrate (βOHB; µmol·L^-1^), and **(D)** the βOHB-to-AcAc ratio at rest and following 30 minutes of exercise. **(E)** A time course of blood β-ketone bodies in mice during sedentary conditions, acute exercise (EX), and refeeding (mmol·L^-1^). **(F)** Area under the curve (AUC) for blood β-ketone bodies during post-exercise refeeding (mmol·min·L^-1^). **(G)** Liver respiratory chain complexes 1-5 (CI-CIV) via immunoblotting. Liver **(H)** phospholipids (PL; µg·mg^-1^), **(I)** diacylglycerides (DAG; µg·mg^-1^), and **(J)** triacylglycerides (TAG; µg·mg^-1^) in sedentary mice. **(K)** Liver TAG (µg·mg^-1^) following 30 minutes of exercise. **(L)** Liver TAG (µg·mg^-1^) in refed mice following exercise. n = 5-9 per genotype. Data are mean ± SEM. *p<0.05 and **p<0.01.

However, TAG accumulated markedly in the livers of KO mice by the end of the 30-minute treadmill bout (Fig. 4K) and remained elevated during refeeding post-exercise (Fig. 4L). Taken together, these results suggest that ketone body homeostasis is shifted towards βOHB and that the suppression of ketone body production in the early stages of refeeding following acute exercise is delayed in mice lacking PCK1. Furthermore, these changes in ketone body metabolism during refeeding post-exercise may be linked to higher liver lipid availability.

### Exercise training diminishes gain in body weight and adiposity in KO mice

Given that KO mice showed dysregulated nutrient metabolism during and following an acute exercise bout, the impact of exercise training on systemic and tissue-specific metabolism was tested in WT and KO mice. Ad libitum body weight, body fat percentage and lean mass percentage were comparable between WT and KO mice prior to exercise training protocols at 12 weeks of age (Fig. 5A-C). Reassessment of these metrics following 6 weeks of exercise training did not show significant differences between genotypes or exercise training protocols (Fig. 5D-F). However, KO-Trained mice gained less body weight and fat mass over 6 weeks of exercise training compared to both sedentary groups (WT-Sed and KO-Sed; Fig. 5G and H). These results suggest that loss of hepatic PCK1 enhanced the protective effect of exercise against diet-induced obesity.

**FIGURE 5.**
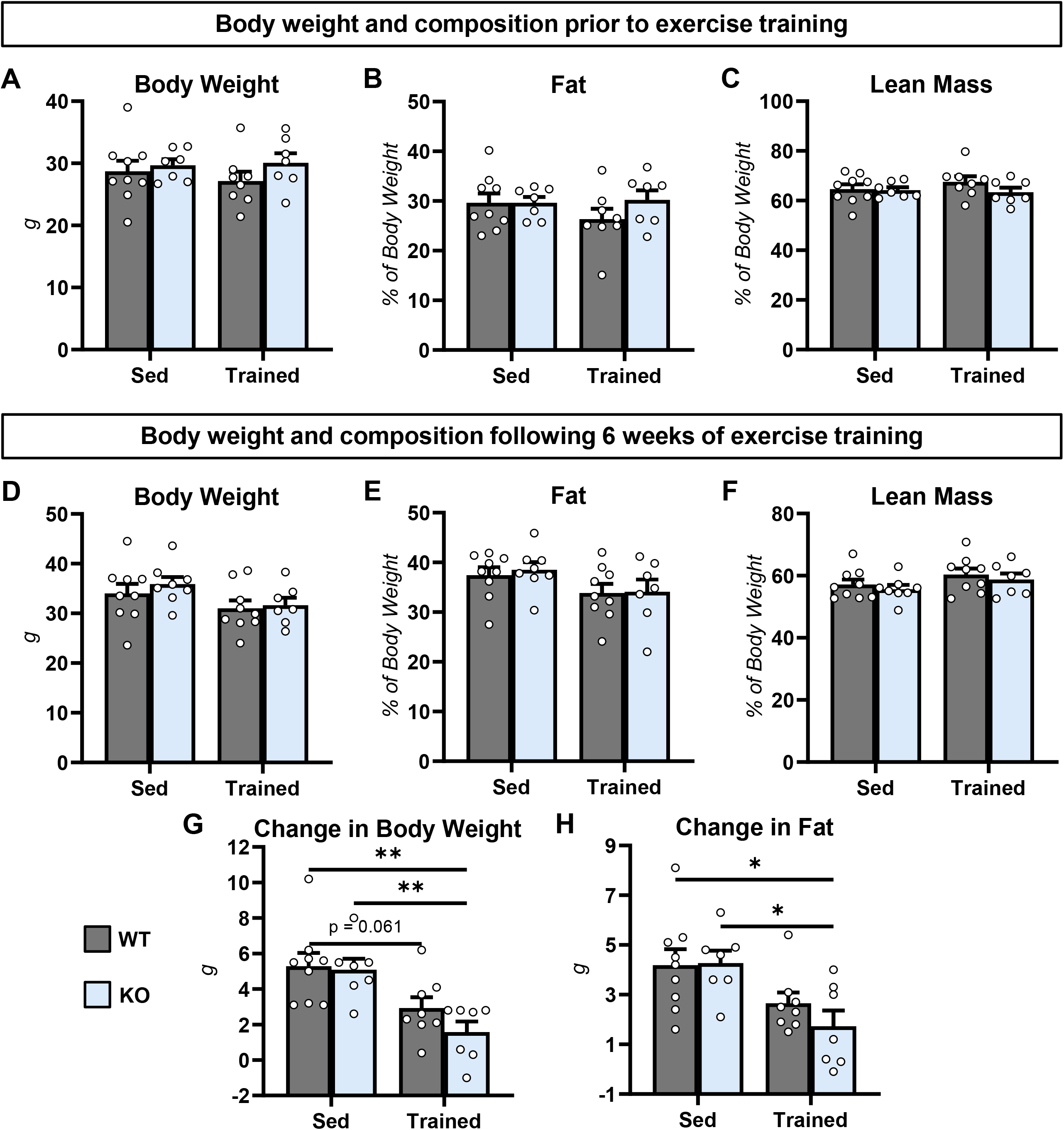
Anthropometric adaptations to exercise training in mice lacking PCK1. Ad libitum fed **(A)** body weight (g), **(B)** fat (% of body weight), and **(C)** lean mass (% of body weight) in 12-week-old mice with a hepatic-specific knockout (KO) of phosphoenolpyruvate carboxykinase 1 and wild type (WT) littermates prior to 6 weeks of exercise training protocols. Ad libitum fed **(D)** body weight (g), **(E)** fat (% of body weight), and **(F)** lean mass (% of body weight) in 18-week-old mice following 6 weeks of exercise training protocols. Change in **(G)** body weight (g) and **(H)** fat mass (g) in mice following the 6 weeks of exercise training protocols n = 7-9 per group. Data are mean ± SEM. *p<0.05 and **p<0.01.

### Exercise training prevents fatty liver in mice lacking hepatic PCK1

Experiments also tested the impact of exercise training on liver metabolism in WT and KO mice (Fig. 6). Liver G6PC and PYGL were decreased in KO-Sed mice compared to WT-Sed mice (Fig. 6A). The decline in these molecular regulators of glycogenolysis and glucose production persisted with training (Fig. 6A). Similar to mice at 12 weeks of age, liver glycogen was diminished in 18-week-old KO-Sed compared to WT-Sed (Fig. 6B). Remarkably, exercise training normalized liver glycogen in KO mice (Fig. 6B). Exercise training also influenced liver lipid metabolism in KO mice. There was no effect on liver PLs (Fig. 6C), but six weeks of treadmill running lowered liver DAGs in KO-Trained compared to KO-Sed mice (Fig. 6D). Liver TAGs were higher in KO mice compared to WT mice under sedentary conditions (Fig. 6E). Unexpectedly, liver TAGs were lower in KO-trained compared to KO-Sed mice (Fig. 6E). The reduction in liver fat could not be explained by altered mitochondrial respiratory chain complexes protein as complexes III (CIII) and complex IV (CIV) were lower in KO mice compared to WT mice in both sedentary and exercise-trained conditions. Ketogenesis is an additional pathway by which the liver can dispose of hepatic fat. Liver ketone bodies were comparable between KO-Sed and WT-Sed mice at 18 weeks of age (Fig. 6G). However, AcAc, βOHB, and the total ketone body (TKB) pool were lower in livers of KO-Trained mice compared WT-Trained mice (Fig. 6G). Liver HMGCS2 protein was similar between all groups and BDH1 was higher in KO mice as indicated by a main effect for genotype (Fig. 6H). These data demonstrate that exercise training prevents liver steatosis in the absence of hepatic PCK1 and that the effect was not be linked to pronounced increases in indices of liver mitochondrial oxidative metabolism.

**FIGURE 6.**
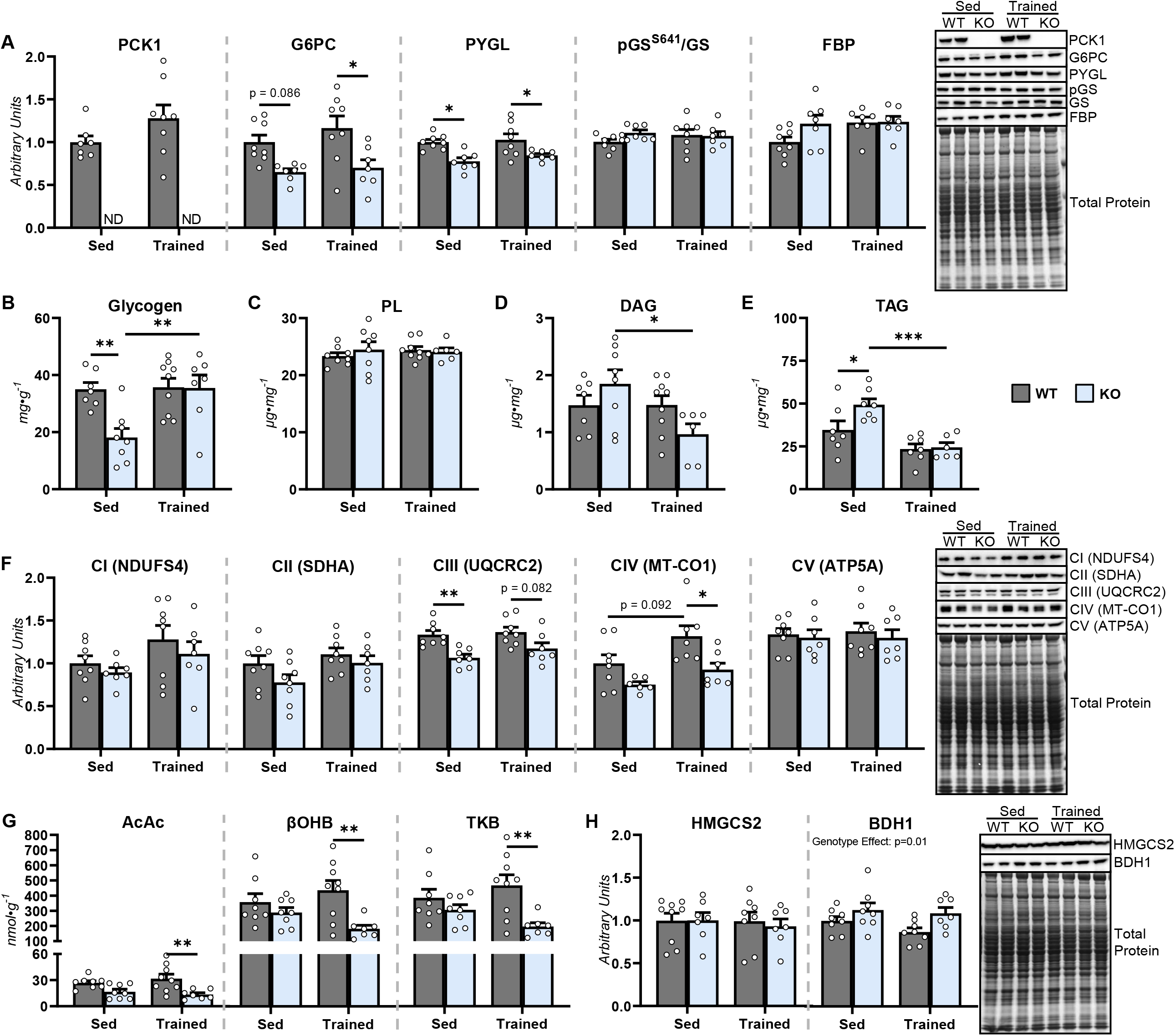
Liver adaptations to exercise training in mice lacking PCK1. Following 6 weeks of exercise training protocols, livers from 18-week-old mice with a hepatic-specific knockout (KO) of phosphoenolpyruvate carboxykinase 1 (PCK1) and wild type (WT) littermates were obtained during fasted, sedentary mice to determine **(A)** liver PCK1, glucose-6-phosphatase catalytic subunit (G6PC), glycogen phosphorylase (PYGL), phosphorylated glycogen synthase-to-total glycogen synthase ratio (pGS^S641^/GS), and fructose 1,6-biphosphatase (FBP) protein via immunoblotting. Liver **(B)** glycogen (mg·mg^-1^), **(C)** phospholipids (PL; µg·mg^-1^), **(D)** diacylglycerides (DAG; µg·mg^-1^), and **(E)** triacylglycerides (TAG; µg·mg^-1^). **(E)** Liver respiratory chain complexes 1-5 (CI-CIV) as determined by immunoblotting and representative immunoblots. **(F)** Liver acetoacetate (AcAc; nmol·g^-1^), β-hydroxybutyrate (βOHB; nmol·g^-1^), and total ketone bodies (TKB; nmol·g^-1^). **(G)** Liver 3-hydroxymethylglutaryl-CoA synthase 2 (HMGCS2) and β-hydroxybutyrate dehydrogenase (BDH1) as determined by immunoblotting and representative immunoblots. n = 6-9 per group. Data are mean ± SEM. *p<0.05, **p<0.01, ***p<0.001. ND, not determined.

### Skeletal muscle adaptations to exercise training are enhanced in KO mice

Given that the liver is an important source of nutrients that provide energy to skeletal muscle during acute exercise, adaptations in muscular performance and metabolism were assessed in WT and KO mice following six weeks of exercise training (Fig. 7). Maximal running speed and cumulative work during an exercise stress test were comparable between all groups of mice prior to initiating exercise training (Fig. 7A and B). In contrast, training tended to improve maximal running speed and work in WT-Trained but not KO-Trained mice (Fig. 7C and D). Nevertheless, mitochondrial respiratory complexes (Fig. 7E) and citrate synthase activity (Fig. 7F) were generally increased in gastrocnemius muscle of KO-Trained. Gastrocnemius SCOT protein was increased in response to exercise compared to sedentary conditions and BDH1 protein was higher in KO mice compared to WT mice as indicated by main effects of exercise and genotypes, respectively (Figure 7G-I). WT-Sed and KO-Sed mice showed comparable muscle glycogen at 18 weeks of age (Fig. 7J). Of note, muscle glycogen was elevated ∼2-fold in KO-Trained mice compared to all other groups (Fig. 7J). Together, these results suggest that loss of liver PCK1 limits improvements in muscular performance but enhances the metabolic adaptations of muscle to exercise training including increased glycogen deposition and mitochondrial oxidative metabolism.

**FIGURE 7.**
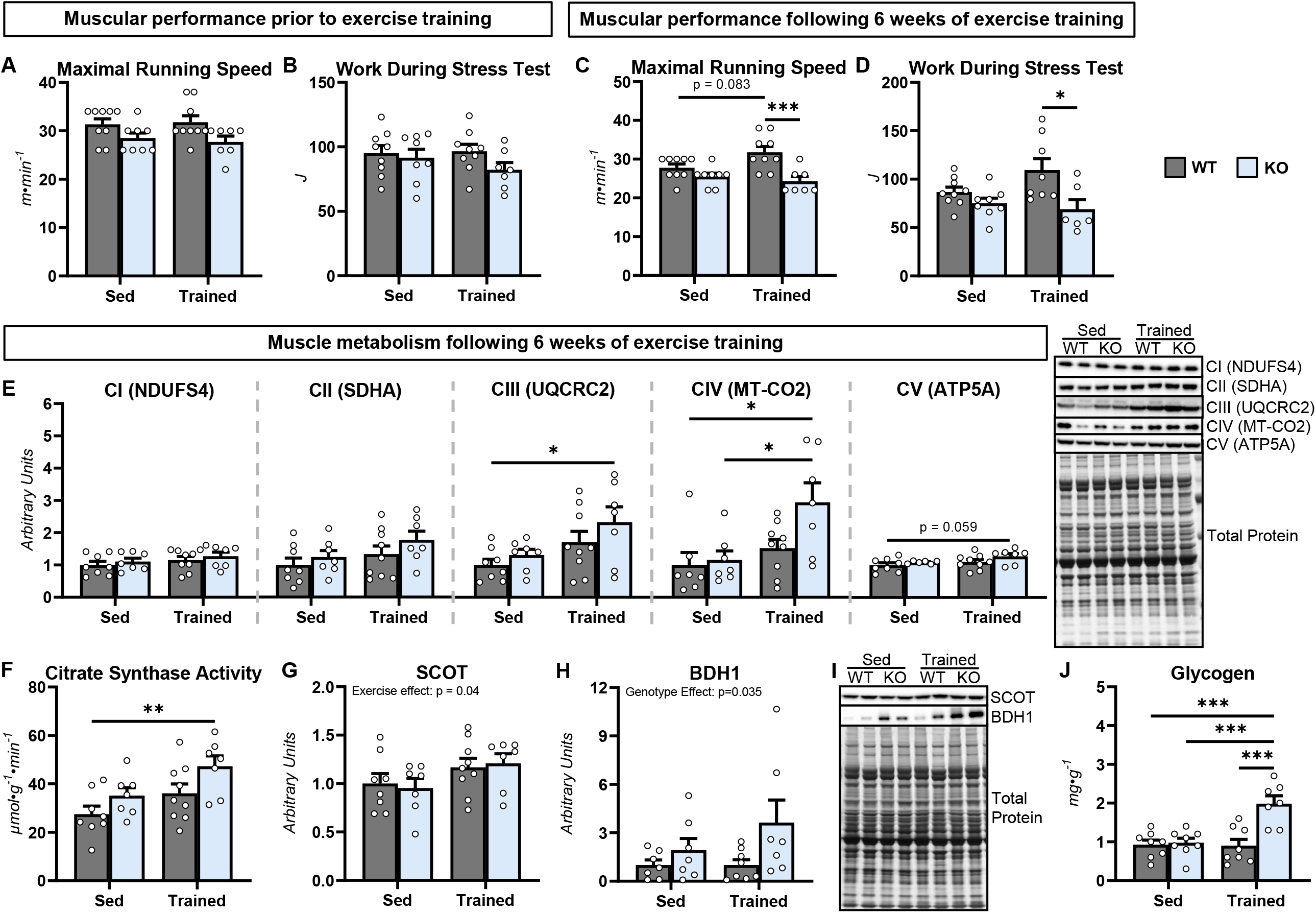
Skeletal muscle adaptations to exercise training in mice lacking PCK1. **(A)** Maximal running speed (m·min^-1^) and **(B)** work (J) during an exercise stress test in 12-week-old mice with a hepatic-specific knockout (KO) of phosphoenolpyruvate carboxykinase 1 and wild type (WT) littermates prior to 6 weeks of exercise training protocols. **(C)** Maximal running speed (m·min^-1^) and **(D)** work (J) during an exercise stress test in 18-week-old mice following 6 weeks of exercise training protocols. Following the 6 weeks of exercise training protocols, gastrocnemius was obtained during fasted, sedentary mice to determine muscle **(E)** respiratory chain complexes 1-5 (CI-CIV) protein via immunoblotting, and **(F)** citrate synthase activity (µmol·g^-1^·min^-1^). Muscle **(G)** succinyl-CoA:3-oxoacid-CoA transferase (SCOT) and **(H)** β-hydroxybutyrate dehydrogenase (BDH1) from fasted, sedentary mice as determined by immunoblotting and **(I)** representative immunoblots. **(J)** Muscle glycogen (mg·mg^-1^) from fasted, sedentary mice. n = 6-9 per group. Data are mean ± SEM. *p<0.05, **p<0.01, ***p<0.001.

## DISCUSSION

Regular exercise is effective at preventing hepatic steatosis (8-12) and in combination with other lifestyle modifications is a first-line intervention for NAFLD (37). The work presented in this study aimed to further define the mechanisms by which exercise training elicits beneficial outcomes in NAFLD. Specifically, we sought to test the hypothesis that repeated bouts of acutely increased hepatic gluconeogenesis are necessary for exercise training to lower liver lipids. The salient findings of this study were i) hepatic PCK1 is required for acute exercise to increase endogenous gluconeogenesis, ii) a single bout of exercise will promote a transient accumulation of liver lipids if gluconeogenesis from phosphoenolpyruvate is not provoked during the muscular work, and iii) repeated provocation of gluconeogenesis from phosphoenolpyruvate is not necessary for exercise training to prevent diet-induced liver steatosis.

Consistent with prior work (20, 21, 23, 24), our experiments showed that KO mice had preserved endogenous glucose production (V_EGP_) and glycemia under fasted, sedentary conditions. Glycogenolysis (V_PYGL_) was similar between genotypes at rest, however, liver glycogen content was decreased in KO mice compared to WT mice in both the fasted and refed states. This was associated with a higher pGS^S641^/GS ratio in livers of KO mice, which could diminish the glycogen synthetic activity of GS (38). These results are consistent with the loss of gluconeogenic capacity, which may cause the mice to transition to the fasted state more rapidly as well as impair the indirect pathway of glycogen synthesis (39) and blunt the repletion of glycogen in the fed state. Endogenous gluconeogenesis from glycerol (V_GK_) did not compensate for loss of PCK1, which is similar to previous findings *in vivo* (21, 24) and isolated perfused liver studies (22, 36). Using isotope labeling and multicompartment flux modeling, Rahim et al. found that the primary compensatory mechanism in mice lacking hepatic PCK1 was an induction of renal gluconeogenesis mediated through increased gluconeogenesis from phosphoenolpyruvate, cataplerosis, and TCA cycle fluxes (24). Since these renal fluxes are encoded in isotope labeling of plasma glucose in KO mice, they likely underlie the blunted but incomplete suppression of *in vivo* gluconeogenesis and cataplerosis and similar TCA cycle fluxes that preserve V_EGP_ and euglycemia in KO mice at rest.

The current study not only confirms previous work but expands on it by quantifying endogenous glucose and oxidative fluxes in KO mice during an acute exercise bout. While WT mice showed a slight increase in blood glucose during exercise, KO mice displayed a progressive decline in circulating glucose during a 30-minute treadmill run. The increase in endogenous glucose production observed in WT mice during acute exercise was impaired in KO mice. This was owing to the inability of muscular work to increase gluconeogenesis (V_Enol_ and V_GK_), cataplerosis (V_PCK_), and TCA cycle fluxes (V_SDH_) in KO mice. The mechanisms responsible for the lack of response in gluconeogenesis from both glycerol and phosphoenolpyruvate to exercise are unclear. Elevated circulating glucagon concentrations and it’s signaling actions are critical in mobilizing liver glycogen and accelerating gluconeogenic flux during exercise (16, 40). While glucagon levels are higher in mice lacking hepatic PCK1 (24), it may be that elevated glucagon stimulates renal gluconeogenesis at rest, however, a further rise in glucagon concentration and/or action is not observed in KO mice during exercise. Notably, the findings from our experiments support the importance of functional coupling between hepatic gluconeogenesis and lipid disposal during acute exercise as liver triacylglycerides were comparable between genotypes at rest, but accumulated in the livers of KO mice during a 30-minute treadmill run.

Our ^2^H/^13^C metabolic flux experiments during acute exercise confirmed mice lacking hepatic PCK1 was a model system in which we could test whether repeated bouts of increased gluconeogenesis are necessary for exercise training to lower liver lipids. Regular exercise concurrent with feeding of a high-fat diet limits the accretion of liver lipids in rodent models (41-44). Consistent with this previous independent work, liver TAG trended lower in WT-Trained mice compared to WT-Sed mice in our study. Twelve weeks of high-fat feeding promoted increased liver TAG in KO-Sed mice compared to WT-Trained mice. Surprisingly, six weeks of treadmill running prevented lipid accumulation in the livers of KO mice. This outcome indicates that hepatic gluconeogenesis from phosphoenolpyruvate and associated oxidative fluxes are not required for exercise training to mitigate diet-induced fatty liver.

Given that exercise training was able to prevent fatty liver in KO mice, further experiments were completed to evaluate the metabolic mechanisms contributing to this beneficial outcome. Hepatic terminal oxidation, as indicated by mitochondrial respiratory chain proteins, was not greater in KO mice undergoing exercise training. Beyond terminal oxidation of fatty acids via TCA cycle and oxidative phosphorylation, the liver can dispose of lipids through ketogenesis (45). Furthermore, ketone bodies are primarily synthesized by the liver and they become an increasingly important fuel source for extrahepatic tissues when glucose availability is limited (46). Prior work found that low-fat fed mice with a global knockdown of PCK1 exhibited a trend towards increased rates of ketogenesis under sedentary, overnight fasted conditions (23). On a high-fat diet, mice with a global PCK1 knockdown had a higher βOHB-to-AcAc ratio but similar ketogenic flux compared to control mice (23). In contrast, mice lacking hepatic PCK1 were previously found to have lower plasma βOHB after a 24-hr fast, but AcAc was not measured (20). In this study, we found a higher circulating βOHB-to-AcAc ratio owing to lower AcAc and comparable βOHB in 12-week-old, untrained, sedentary KO mice compared to WT mice. At 18-weeks of age, KO-Sed mice had similar liver AcAc and βOHB compared to WT-Sed mice. Together with the existing literature, our results suggest that ketogenesis is not appreciably impacted by loss of hepatic PCK1 under sedentary, fasting conditions. The absence of pronounced changes in indices of ketogenesis is consistent with the preserved euglycemia in sedentary KO mice.

Very little is known about how exercise affects ketogenesis in mice. A considerable limitation has been the inability to acquire serial samples to measure blood ketone bodies in running mice, which we overcome here with the implantation of an arterial catheter. Circulating βOHB declined with the onset of exercise, but returned to resting levels by 30-minutes of exercise in both WT and KO mice. In addition, the magnitude of post-exercise ketosis was comparable between genotypes. Interestingly, the decline in βOHB availability early in an exercise bout as well as the post-exercise ketosis in mice is very similar to that observed in fasting humans (47-50). This is attributed to ketone body disposal being elevated quickly during muscular work while the rise in ketogenic flux during exercise is delayed and outpaces ketone body disposal upon cessation of exercise (47-50). Notably, arterial βOHB decreased less rapidly upon refeeding following exercise in KO mice compared to WT mice. The factors responsible for the higher ketone bodies in KO mice during refeeding were not determined, however, this ketone body phenotype is congruent with higher rates of ketogenesis during conditions of low glucose/glycogen and higher lipid availability (51). Furthermore, the prolonged rise of ketogenesis during refeeding after an acute exercise bout may aid in diminishing the accumulation of liver lipids in KO mice. Future experiments testing the impact of ketogenic insufficiency on the ability of exercise to prevent hepatic steatosis are warranted.

In addition to the disposal of lipids in hepatic metabolic pathways, extrahepatic metabolism was also considered in the prevention of fatty liver in exercise-trained KO mice. Repeated bouts of muscular work stimulate metabolic adaptations in skeletal muscle to improve nutrient homeostasis and facilitate a more efficient provision of ATP during future exercise bouts (52). These adaptations include greater glycogen storage and increased mitochondrial function, number, and quality control (53, 54). Here we showed that exercise training in KO mice led to a more pronounced increase in skeletal muscle glycogen deposition, citrate synthase activity, and mitochondrial respiratory chain proteins. These results suggest that loss of hepatic PCK1 enhances exercise-induced adaptations in skeletal muscle mitochondrial oxidative metabolism. Enhanced mitochondrial metabolism could elevate skeletal muscle lipid disposal and, subsequently, lessen fatty acid delivery to the liver. Findings in our study are in-line with this hypothesis as exercise training limited the gain in adiposity in KO mice to a greater extent than WT mice. In addition, liver AcAc and βOHB were lower in KO-Trained mice. Given that fatty acids are a potent stimulator of hepatic ketogenesis (23, 51, 55), the reduction in ketone bodies may be due to reduced supply of lipids to the liver.

The stimuli driving enhanced mitochondrial adaptations to exercise training in the skeletal muscle of KO mice are not currently known. In speculation, the lower glucose availability (circulating glucose and skeletal muscle glycogen) during and following muscular work is an important factor. Hansen et al. tested metabolic adaptations in individuals completing a 10-week knee extensor training protocol under conditions of low and high muscle glycogen (56). Exercise undertaken with both low and high glycogen elevated skeletal muscle citrate synthase activity, however, the rise was more pronounced under conditions of the low glycogen (56). In agreement, follow-up studies have routinely reported superior increases in skeletal muscle mitochondrial respiratory chain proteins, TCA cycle and β-oxidation enzyme activity, and fat oxidation in response to low glycogen exercise training (57-60). Further support for the role of glucose availability in mitochondrial adaptations to exercise comes from studies evaluating the impact of hyperglycemia. An increasing number of studies have linked impaired fasting glucose to blunted skeletal muscle adaptations to exercise training (61, 62). In particular, Osler et al. reported that in individuals with impaired glucose tolerance, those with impaired fasting glucose did not show an increase in markers of muscle mitochondrial respiratory function following exercise training (63). As such, future studies designed to maintain euglycemia in KO mice during acute exercise may help to clarify the importance of hepatic gluconeogenesis in mediating skeletal muscle adaptations to exercise training and the role of these adaptations in preventing hepatic steatosis.

In summary, this study shows that repeated bouts of acutely increased gluconeogenesis from phosphoenolpyruvate are not required for exercise training to prevent diet-induced liver lipid accretion. Our results suggest that the efficacy of exercise training to lower lipids in the absence of PCK1 is linked to a delayed suppression of ketogenesis upon refeeding following acute exercise as well as enhanced metabolic adaptations in skeletal muscle precipitated by hypoglycemic episodes during each acute exercise bout.

## ACKNOWLEDGEMENTS

The authors thank the University of Minnesota Molecular Medicine Metabolomics Core for completing liver and plasma ketone body measurements, the Masonic Cancer Center Analytical Biochemistry shared resource at the University of Minnesota (NIH P30 CA77598) for access to GC–MS instrumentation, and the Vanderbilt University Medical Center Lipid Core (NIH DK059637 and DK020593) for quantifying liver lipids. Figure schematics were created using BioRender.com.

## AUTHOR CONTRIBUTIONS

CCH conceived and designed the experiments. FIR, GLS, WCV, DBS, and CCH performed experiments. FIR and CCH analyzed and interpreted data. FIR and CCH drafted the manuscript. All authors edited and revised the manuscript. All authors approved the final version of the manuscript for publication.

